# Interaction between perineuronal nets and ketamine in antidepressant action

**DOI:** 10.1101/2021.05.30.446326

**Authors:** Calvin K. Young, Kachina G. Kinley, Neil McNaughton

## Abstract

Depression is highly prevalent, increases suicide risk, and is now the leading cause of disability worldwide. Our ability to treat depression is hampered by the lack of understanding of its biological underpinnings and of the mode of action of effective treatments. We hypothesised that the scaffolding proteins in the medial frontal cortex play a major role in effective antidepressant action. We implanted cannulae into the infralimbic cortex to inject chABC and locally remove perineuronal nets and then tested for antidepressant effects with the forced swim test. We further tested if systemic injections of ketamine had an additive effect. Our preliminary data indicate that neither the removal of these scaffolding proteins nor ketamine were sufficient to decrease depression-like behaviour, but may interact synergistically to decrease immobility time in the forced swim test.

## Introduction

Depression is on the rise worldwide (WHO 2017). This trend warrants attention as depression can significantly decrease quality of life, is associated with poorer prognosis of other conditions, and places heavy demands on the healthcare system (Roy-Byrne, Stang et al. 2000, Oakley-Browne, Wells et al. 2006, Brakowski, Spinelli et al. 2017). There are various treatments for depression, including antidepressant medication and cognitive behavioural therapy. However, we know little about the biological basis of the disorder and about how effective treatments work.

Current first-line treatment is through modulation of the monoamine system. The effectiveness of monoamine treatments, particularly commonly used selective serotonin reuptake inhibitors (SSRIs) such as Prozac, suggests that patients with depression have reduced levels of serotonin, dopamine, and/or norepinephrine (Hirschfeld 2000). However, alternative forms of treatment including prefrontal cortex deep brain stimulation, electric shock therapy, and cognitive behaviour therapy have proven to be effective, even though they do not appear to act directly on the monoamine system. One thing all these treatments have in common is that they increase synaptic plasticity by increasing brain-derived neurotrophic factor – BDNF (Nibuya, Morinobu et al. 1995, Gersner, Toth et al. 2010, Duman and Aghajanian 2012). Thus, the aetiology of depression and its effective treatment may hinge on quantifiable wiring abnormalities in the brain.

A growing literature implicates perineuronal nets (PNN) in psychiatric conditions (Berretta 2012). PNNs are organised structures composed of chondroitin sulfate proteoglycans (CSPG) and extracellular matrix (Deepa, Carulli et al. 2006), which is crucial in stabilising neural connections (Bahia, Houzel et al. 2008). Degradation of GSPGs by the enzyme chondroitinase ABC (chABC) has been shown to assist in recovery from spinal cord injury (Bradbury, Moon et al. 2002), promote the erasure of traumatic memories (Gogolla, Caroni et al. 2009), and reverse developmental critical periods (Pizzorusso, Medini et al. 2002).

SSRIs (the most commonly prescribed antidepressant medication) have demonstrated effects on PNNs (Maya Vetencourt, Sale et al. 2008, Karpova, Pickenhagen et al. 2011, Scali, Begenisic et al. 2013); producing reduced PNN staining in the mPFC, which suggests that PNNs are altered by the antidepressant effects of the treatment (Ohira, Takeuchi et al. 2013).

Ketamine has long-lasting antidepressant effects (Browne and Lucki 2013) that, unlike SSRIs, are immediate. Ketamine is also effective in those resistant to other antidepressant treatments (Gill, Gill et al. 2021). However, it has not been tested whether or not ketamine modifies PNNs in the mPFC relating to depression-like behaviours. Here we tested whether the degradation of PNNs underlies antidepressant treatment by allowing greater freedom for synaptic re-wiring. We probe how systemic ketamine changes both PNNs and depression-like behaviours in rodents. Additionally, we examine whether the direct, non-pharmacological removal of PNNs in the mPFC using chABC results in antidepressant behavioural effects.

## Methods

Sixteen male Wistar rats (350-430 g) obtained from the Department of Laboratory Animal Sciences at the University of Otago received bilateral guide cannula implantation targeting the infralimbic cortex (IL; AP: +3.2 mm; ML: +/-1.4 mm @ 10°; DV: -4.7 mm) after seven days of acclimatisation. After a seven-day recovery period, each rat was subjected to a 15-minute swim session as part of the forced swim test (Slattery and Cryan 2012). Thirty minutes after the swim, rats were first injected with 0.5 μl control enzyme (penicillinase, P0389-1KU, Sigma, USA) or chABC (C3667-10UN, Sigma, USA) through an injection cannula inserted to the guides (i.e. into the IL) at a rate of ∼0.3 μl/min. The injection cannulae were left in place for 5 minutes after the injection. Injections were always made first in the right hemisphere, then the left. Immediately after this, 10 mg/kg of ketamine (ket) or volume-matched physiological saline (sal; 0.9%) was injected subcutaneously. This yielded a 2×2 design with 4 rats in each group receiving a combination of ketamine/saline (ket/sal) and penicillinase/chABC (pen/chABC) injections. The rats were monitored for at least 30 minutes after the last injection. Twenty-three hours later, rats were exposed to another five-minute swim session. Thirty minutes after the second exposure, rats were transcardially perfused with phosphate-buffered saline followed by 4% paraformaldehyde (PFA, in 0.1 M phosphate buffer). The extracted brains were further fixed in 4% PFA for 24 hours, transferred to a 30% sucrose (in phosphate-buffered saline) solution for cryoprotection prior to sectioning. Forty-micron sections were made in a cryostat and stored in Tris-buffered saline (TBS) prior to immunostaining.

Immunostaining of PNNs and parvalbumin positive interneurons (PV+) were visualised by WFA (*Wisteria floribunda agglutinin*) conjugated fluorescein (VEFL1351, Vector Laboratories, USA) and rabbit anti-parvalbumin (PV24, Swant, Switzerland) antibody. Sections were first washed with TBS, followed by incubation in a blocking solution (3% normal goat serum; NGS, and 0.03% Triton-X) to reduce background staining and increase the permeability of the tissue. WFA-fluorescein (1:1000) and rabbit anti-PV (1:1000) were added to TBS+3% NGS to incubate overnight. Additional washes in TBS were made before incubation with a goat anti-rabbit Cy3 secondary antibody (1:200; 111-165-144 1:200, Jackson ImmunoResearch, USA) for 4 hours. Final TBS washes were made before sections were mounted on pre-cleaned glass slides and coverslipped with anti-fade glue (F6057, Sigma, USA) with DAPI (a nuclear stain).

Microscope images were acquired with a Nikon (C2plus) confocal microscope at 20x magnification. Two laser lines (488 nm and 561 nm) were used to excite WFA-fluoroscein and PV (Cy3), respectively. We centred our field of view aimed at the prelimbic cortex (PrL, immediately dorsal to IL) and collected a montage (between 7×9 to 9×13 grids, at 512×512 pixels per grid) with a stack of 9 images in 1 micron steps in the z-axis. The acquired images were imported to ImageJ/FIJI (https://fiji.sc, NIH, USA) for further analysis. Firstly, the pseudo-coloured images were pre-processed into 8-bit greyscale images. Regions of interest (ROI) were drawn around IL where this was free of damage caused by the cannulae. Four ROIs (two from each hemisphere) were exported in this fashion for each rat. These images were submitted to semi-automated analysis using PIPSQUEAK (Slaker, Harkness et al. 2016). Thresholds of detection were manually adjusted to include most, if not all of the signal in each channel (i.e. WFA and PV). Auto-fluorescence was manually removed, and missed positive staining was added before single-cell ROIs were processed for single/double labelling analysis.

Statistical analyses were carried out in SPSS (IBM, USA). Data transformation, curve fitting, and figure preparation were performed in Matlab (MathWorks, USA). For the FST, the number of right hind leg kicks per second was taken as the quantitative measure for swimming activity (Warden, Selimbeyoglu et al. 2012). Particularly, a distinction was made between full kicks and half kicks, with the latter more prominent in attempts to stay afloat. This measure was averaged in 5 second periods to reduce variability, and any bin returning less or equal to 1 was counted as immobile. In addition, immobility latency was calculated as the midpoint of a sigmoidal fit of the “kick-frequency” data over time. The swim-to-float curves of four rats’ (2 from the ket/pen group, and 2 from the sal/pen group) did not conform with a sigmoidal curve, and the midpoint of a logarithmic function was used in these cases instead. Histological results were exported as a text file, which included the number of cells detected and background-subtracted intensity used here for further analysis. For both datasets, two-way analyses of variance (ANOVA) were carried out. Pearson correlation coefficients are reported for the relationship between behaviour and histological findings. One rat (ket/chABC group) died due to surgical complications and another rat (sal/pen group) had unusable brain sections due to high background fluorescence; the latter rat contributed to FST data but not histological data.

## Results

First, we demonstrated that the cannula tips were located within the IL region (e.g. Figure 1A). Cannula tips from only one rat from the ket+pen group could not be located (damage to the frontal cortex by dura during brain extraction) but were presumed to be in the PrL/IL region from available histology. Figure 1A also shows strong colocalization (yellow) between WFA (green) and PV (red); in contrast to chABC-digested tissue (Figure 1B) where darker patches, as well as a red (PV-only) signal can be seen around the injected area and along the guide cannula tracks. At 60x magnification (Figure 1C), colocalization was clear with most of the PV neurons WFA+. A zoomed-in example of chABC-treated tissue (Figure 1D) is shown with a white line placed along the border of the PNN digest. The left side is more ventral to the cannula tip, and WFA staining appears to be unaltered. In contrast, the right side is immediately below the cannula track and clearly lacks WFA (green) staining in the presence of PV (red).

**Figure 1.**
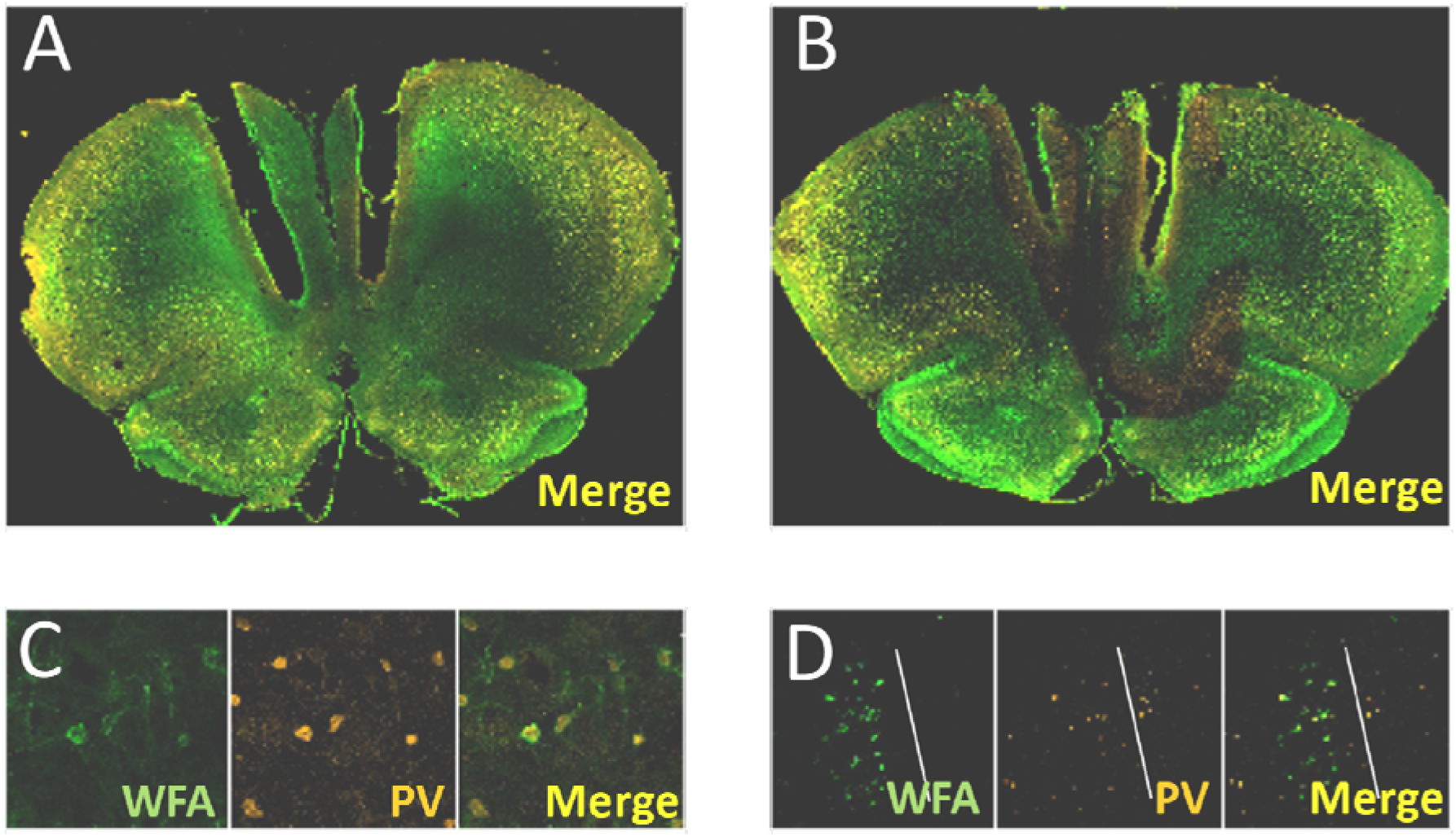
PNN digest with chABC with WFA and PV staining. **A:** An example of a vehicle-injected prefrontal area with WFA and PV staining intact compared to **B:** chABC injected tissue, where clear elimination of WFA signal around the cannula track can be seen. **C:** 20x magnification of representative co-staining of WFA and PV, and **D:** zoomed in view of PV staining without WFA at the “border” (white line) of chABC digest.

The amount of immobility in the FST was highly variable. Although we did observe the lowest variance and mean value in the ket/chABC condition (Figure 2A), there was no interaction between systemic (i.e. sal/ket) and intracerebral (pen/chABC) administration (F_(1, 10)_ = 2.299, *p* = .16), or any effect of chABC or ketamine on their own (F_(1, 10)_ = .997, *p* = .34 and F_(1, 10)_ = 1.150, *p* = .31, respectively). Although variance again was low for the ket/chABC group (Figure 2B), there is no effect of ketamine, chABC, or a combination of the treatments (F_(1, 10)_ = .008, *p* = .93, F_(1, 10)_ = 1.354, *p* = .27, F_(1, 10)_ = .623, *p* = .45, respectively).

**Figure 2.**
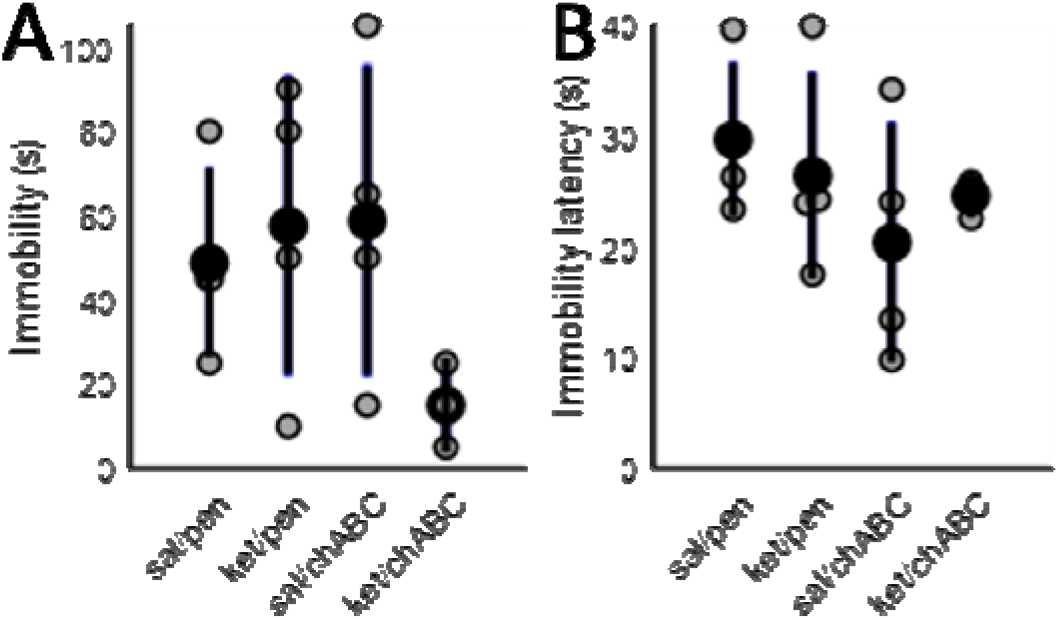
Immobility measures in the FST. **A:** Mean and SE (black) for the time spent immobile in the FST and; **B:** latency to first bout of immobility.

As expected, none of our treatments altered the expression of PV, with the number of PV+ neurons (Figure 3A), total area of PV signal (Figure 3B) and background-corrected intensity (Figure 3C) all comparable (all tests are non-significant, *p* > .13). Digestion of PNNs with chABC did decrease the association of PV+ neurons with WFA (Figure 3D; F_(1, 10)_ = 16.017, *p* < .005) with no effect of ketamine (F_(1, 10)_ = .33, *p* = .86) or the combination of chABC with ketamine (F_(1, 10)_ = .002, *p* = 962).

**Figure 3.**
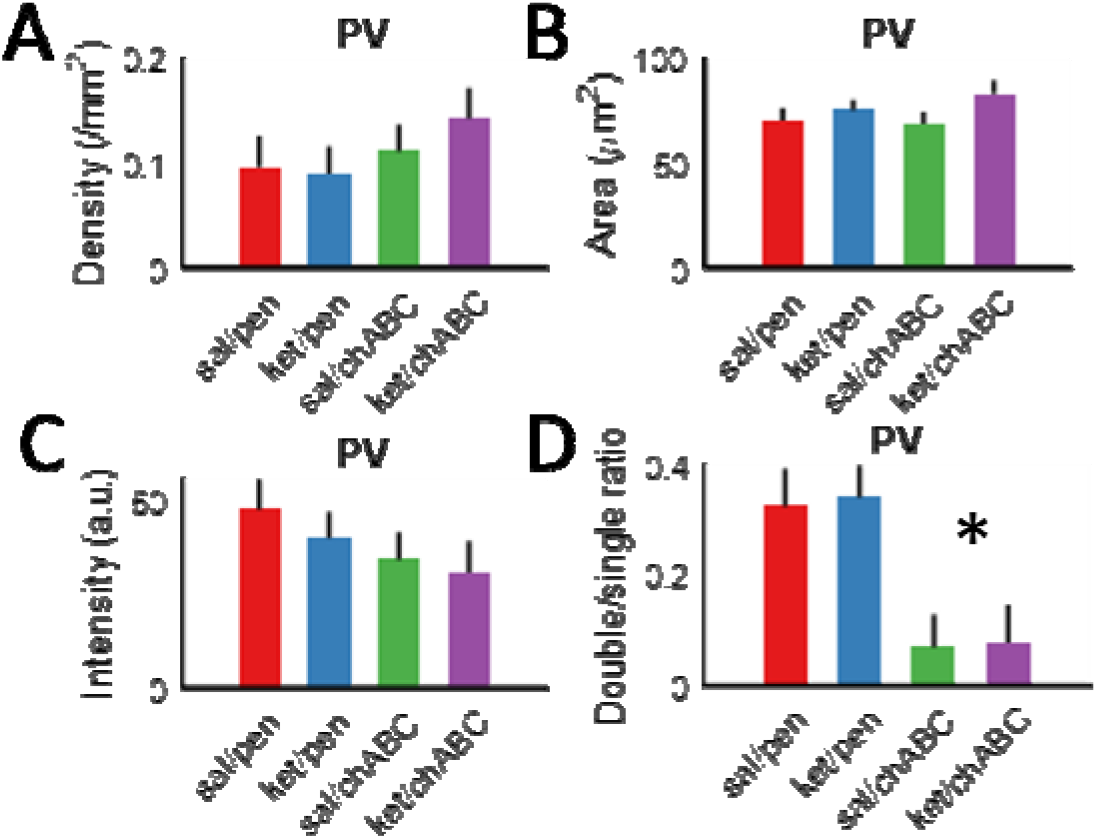
Characteristics of PV staining in the IL. **A:** PV cell count density, **B:** total area (sum of cell sizes), **C:** fluorescence intensity, and **D:** ratio of PV (single) and PV+WFA (double) stained cell counts.

In contrast, chABC greatly reduced the density of WFA staining (Figure 4A; F_(1, 10)_ = 22.021, *p* < .001) with no interaction with ketamine injection (F_(1, 10)_ = .59, *p* = .46). There is no effect of ketamine on the density of WFA staining (F_(1, 10)_ = 391, *p* = .55). The average area of counted WFA staining was not different for chABC, ketamine or a combination of treatments (Figure 4B; all tests non-significant, *p* > .29). Mirroring changes seen in WFA density, the background-corrected intensity of the WFA signal was decreased after chABC treatment (Figure4C; F_(1, 10)_ = 12,711, *p* < .005) but not after ketamine (F_(1, 10)_ = 1.307, *p* =.28), with no interaction between the two treatments F_(1, 10)_ = .253, *p* = .63). There was no difference in the ratio of WFA+ neurons that were also PV+ (Figure 4D; all tests non-significant, *p* > .27).

**Figure 4.**
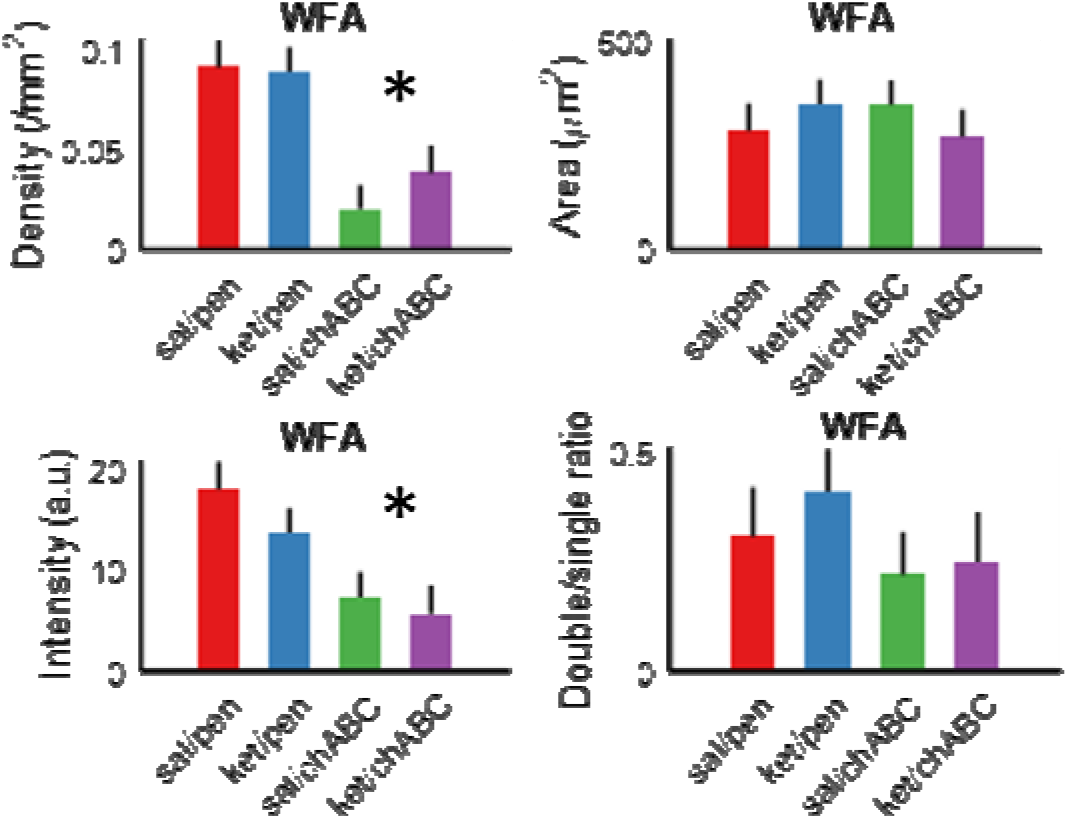
Characteristics of WFA staining in the IL. **A:** WFA cell count density, **B:** total area (sum of cell sizes), **C:** fluorescence intensity, and **D:** ratio of WFA (single) and WFA+PV (double) stained cell counts.

**Figure 5.**
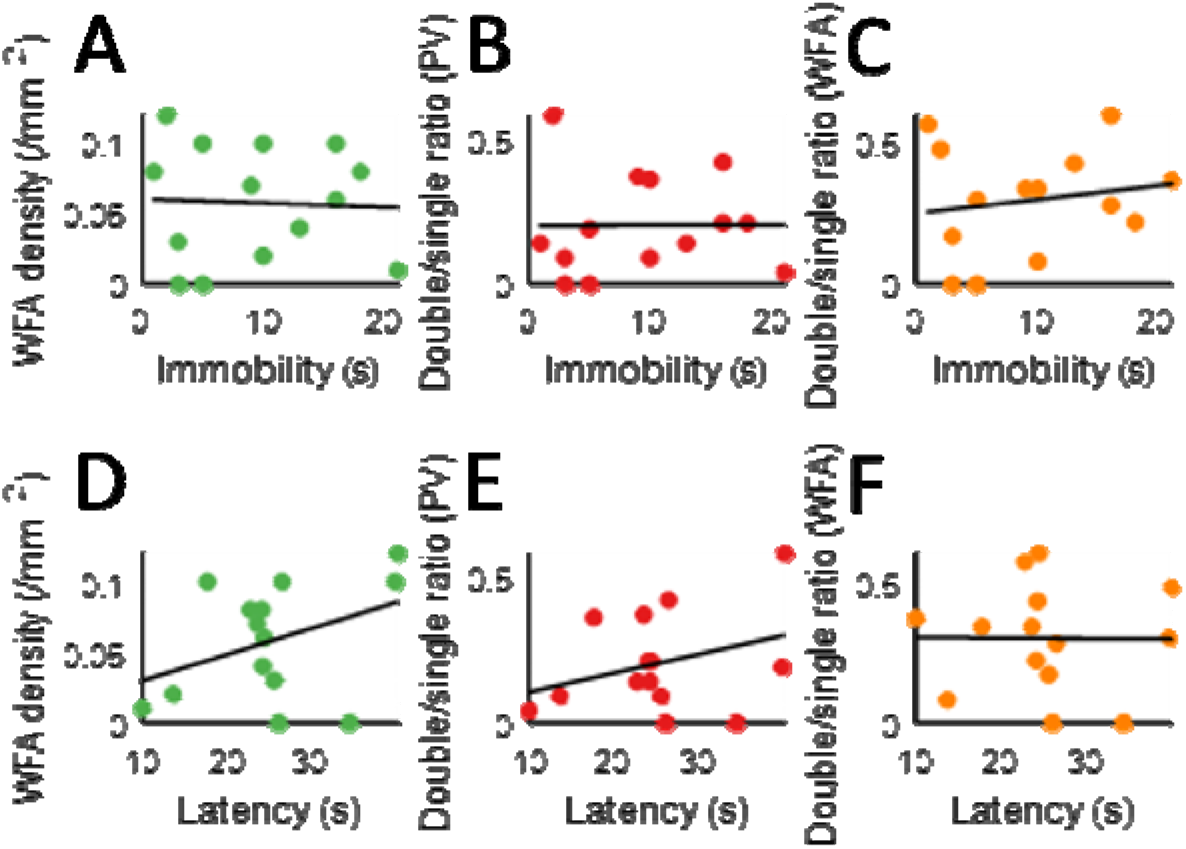
Relationship between WFA/PV staining and immobility in the FST. Correlaitons between **A:** WFA density, **B:** PV+WFA/PV labelling ratio, and **C:** PV+WFA/PV labelling ratio and time spent immobile in the FST. **D-F:** Same histological measures correlated against the latency to first bout of immobility in the FST.

Although there was a clear histological outcome for chABC treatment, the behavioural outcome was inconclusive. To account for potential differences in the extent of chABC digest in relation to behaviour, we carried out correlation analysis in an attempt to establish possible dependencies. We found no relationship between the time spent immobile in the FST and: 1) the density of WFA staining (Pearson’s *r*(14) = -.04, *p* = .89); 2) the proportion of PV+ that are also WFA+ (Pearson’s *r*(14) = .01, *p* = .98) or; 3) the proportion of WFA+ neurons that are PV+ (Pearson’s *r*(14) = .16, *p* = .58). Although correlations of the same histological measurements with immobility latency was more consistent with the hypothesised relationship, there was no detected significant correlation with: 1) density of WFA staining (Pearson’s *r*(14) = .38, *p* = .18); 2) the proportion of PV+ that are also WFA+ (Pearson’s *r*(14) = .32, *p* = .26) or; 3) the portion of WFA+ neurons that are PV+ (Pearson’s *r*(14) = -.01, *p* = .98).

## Discussion

In the current study, we sought to establish a direct link between the integrity of PNNs and depression-like behaviour in the FST. Our data indicate that 10mg/kg ketamine was not effective as an antidepressant in naïve rats tested with FST, and did not change IL WFA staining. We did not observe any antidepressant effect of PNN removal in the IL and surrounding areas, despite an almost complete elimination of PNNs.

We were not able to replicate various studies showing ketamine having an antidepressant effect at sub-anaesthetic doses (i.e. 10 mg/kg) injected 24 hours prior to the test (Fukumoto, Toki et al. 2017, Browne, Falcon et al. 2018, Wan, Feng et al. 2018). We were also not able to demonstrate a clear effect of chABC on depression-like behaviour in the FST. However, we do note that ketamine+chABC treatment did seem to yield consistently favourable results in relation to our hypothesis but the effect was not found to be statistically significant. One obvious weakness of the current study is the low number of samples in each condition. Previous studies have used a minimum of 8+ animals (Fukumoto, Toki et al. 2017, Browne, Falcon et al. 2018, Wan, Feng et al. 2018) to obtain statistically significant differences between control and experimental groups. Our study is somewhat underpowered to sufficiently answer our research questions, but additional data is currently being analysed to clarify the findings described here. We note recent studies have highlighted the relevance of stress levels in rodents relating to ketamine’s antidepressant effects. Specifically, ketamine increased axiogenic behaviours and decreased social interaction time as would stress-induced depression models in naïve Wistar rats (Silvestre, Nadal et al. 1997). In mice, ketamine produced antidepressant effects in stressed mice but had the opposite effect in non-stressed mice (Fitzgerald, Yen et al. 2019, Trofimiuk, Wielgat et al. 2019), an effect also observed in humans (Nugent, Ballard et al. 2019). Despite high variance, our behavioural data indicated an overall pro-depressant effect as measured by immobility in FST. An expanded behavioural battery and the manipulation of pre-treatment stress levels will shed more light on the complex interaction between stress, treatment outcome and the role of structural plasticity.

Traditionally, FST scoring involves the scorer assessing the “dominant” behaviour in 5 s bins to provide a single binary assessment (i.e. immobility or otherwise) of animals’ activity in the 5 min test (Slattery and Cryan 2012). As an attempt to move away from subjective measures of FST behaviour, we employed a more objective measure of hind leg kicks in FST (Warden, Selimbeyoglu et al. 2012). However, we note that our division of full kicks (full extension of the hind leg) and half kicks (less than full extension of the hind leg) does involve a certain degree of subjectivity. Regardless, it is important to point out that we did not score the FST in a conventional way, thus it is unclear if the lack of effect may be due to scoring differences. In addition, we did not differentiate “climbing” (vertical movement) and “swimming” (horizontal movement) behaviours in our current attempt. We are currently applying alternative algorithms that can automatically score behaviour more objectively (Yuman, Idaku et al. 2008), and/or use deep learning algorithms (Mathis, Mamidanna et al. 2018) to carry out more detailed kinematic analyses of FST behaviour.

It is unclear what effect, if any, the damage as a result of cannula implant/intracerebral injection has on PNN/PV expression and FST behaviour. We targeted the IL due to its apparent central role in depression and antidepressant action (Covington, Lobo et al. 2010, Scopinho, Scopinho et al. 2010, Slattery, Neumann et al. 2011, Seese, Chen et al. 2013, Chang, Chen et al. 2014, Fuchikami, Thomas et al. 2015). IL inactivation and lesion have produced opposite effects on FST immobility in past studies (Scopinho, Scopinho et al. 2010, Slattery, Neumann et al. 2011, Chang, Chen et al. 2014). Our short implant to testing time window (5-7 days), as well as potential additional damage done by insertion/removal of the dummy and injection cannulae 24 hours before proper FST testing has not been adequately controlled for. Additional data are needed to account for the potential acute effects of lesion/injections into the IL area on FST behaviour. In addition to potential issues lesions impose on behaviour, their effects on histological analysis are evident. Although we were able to obtain data from the IL by using sections that immediately surrounding the damaged tissue (where injections were made) it is clear that the damaged area is not uniform across rats and thus creates variation in where exactly viable histological analyses could be carried out. In future studies, PNN modification relating to ketamine may have to be done independently to avoid confounds caused by local lesions. However, similar issues relating to localised intracerebral PNN digest cannot be solved in the same way. The use of biocompatible hydrogels (Clarkson, Parker et al. 2015) for slow release or a combination of transgenic/viral vector techniques (Gossen and Bujard 1992, Madisen, Zwingman et al. 2010, Zhao, Muir et al. 2011) may be used to circumvent the effects of lesions relating to the use of implanted cannulae described here.

The hippocampus and the amydgala, two areas that have been found to undergo activity changes in depression (Zeng, Shen et al. 2012) may reveal further insights to the effectiveness of our manipulations without the confounds described in IL above due to tissue damage.

To summarise, we failed to replicate the therapeutic effects of ketamine on depression-like behaviour in the FST. Consistent with the lack of effect, we found no evidence of PNN modification by systemic ketamine injections. Our hypothesis of PNN removal being antidepressant is not fully supported; only a combination of systemic ketamine and IL chABC injection resulted in a trend to an antidepressant effect, despite dramatic removal of PNNs by chABC in the IL and surrounding regions. Together, our data provide preliminary clues on how ketamine and chABC may act synergistically to produce antidepressant effects. Additional experiments are currently underway to provide adequate statistical power for firmer conclusions.

## Acknowledgements

KGK was supported by a summer research scholarship by the Neurological Foundation of New Zealand. The project was supported by a University of Otago Research Grant.

## Contributions

CKY conceived the study, carried out experiments, confocal image acquisition and data analyses. KGK carried out behavioural scoring, immunohistochemistry, confocal image acquisition and image analysis. NM provided supervision and secured funding for the research. CKY prepared the manuscript and all authors reviewed and edited the manuscript.

